# Abiotic and biotic drivers underly short- and long-term soil respiration responses to experimental warming in a dryland ecosystem

**DOI:** 10.1101/2020.01.13.903880

**Authors:** Marina Dacal, Pablo García-Palacios, Sergio Asensio, Beatriz Gozalo, Victoria Ochoa, Fernando T. Maestre

**Affiliations:** Departamento de Biología y Geología, Física y Química Inorgánica, Universidad Rey Juan Carlos, C/ Tulipán s/n, 28933 Móstoles, Spain; Instituto Multidisciplinar para el Estudio del Medio “Ramon Margalef”, Universidad de Alicante, Carretera de San Vicente del Raspeig s/n, 03690 San Vicente del Raspeig, Spain; Departamento de Ecología, Universidad de Alicante, Carretera de San Vicente del Raspeig s/n, 03690 San Vicente del Raspeig, Spain

**Keywords:** soil respiration, biocrusts, dryland, microbial thermal acclimation, short-term vs long-term warming, soil temperature, soil moisture

## Abstract

Soil carbon losses to the atmosphere through soil respiration are expected to rise with ongoing temperature increases, but available evidence from mesic biomes suggests that such response disappears after a few years of experimental warming. However, there is lack of empirical basis for these temporal dynamics in soil respiration responses, and of the mechanisms underlying them, in drylands, which collectively form the largest biome on Earth and store 32% of the global soil organic carbon pool. We coupled data from a ten-year warming experiment in a biocrust-dominated dryland ecosystem with laboratory incubations to confront 0-2 years (short-term hereafter) vs. 8-10 years (long-term hereafter) soil respiration responses to warming. Our results showed that increased soil respiration rates with short-term warming observed in areas with high biocrust cover returned to control levels in the long-term. Warming-induced increases in soil temperature were the main driver of the short-term soil respiration responses, whereas long-term soil respiration responses to warming were primarily driven by thermal acclimation and warming-induced reductions in biocrust cover. Our results highlight the importance of evaluating short and long-term soil respiration responses to warming as a mean to reduce the uncertainty in predicting the soil carbon – climate feedback in drylands.

## Introduction

Soil respiration is expected to increase with global warming (Davidson & Janssens, 2006; Kirschbaum, 2006), contributing to enhance atmospheric CO_2_ concentration. Thus, warming-induced soil carbon (C) losses via soil respiration may lead to a positive C cycle−climate feedback (Tucker, Bell, Pendall, & Ogle, 2013), which is indeed embedded into the climatic models of the IPCC (Ciais et al., 2014). However, most experiments conducted to date on this topic have typically lasted less than four years (Wang et al., 2014), and there is growing evidence showing that elevated soil respiration rates may gradually be offset towards ambient values after a few years of experimental warming (Kirschbaum, 2004; Luo, Wan, Hui, & Wallace, 2001; Melillo et al., 2017, 2002). Multiple mechanisms have been hypothesized to explain such transient effects of warming on soil respiration. For instance, the thermal acclimation of soil microorganisms to the ambient temperature regime (Bradford et al., 2019; Dacal, Bradford, Plaza, Maestre, & García-Palacios, 2019) and the depletion of labile soil C sources (Hartley, Hopkins, Garnett, Sommerkorn, & Wookey, 2008; Schindlbacher, Schnecker, Takriti, Borken, & Wanek, 2015) may drive soil respiration responses to warming over time. Additionally, and given that soil temperature and moisture are the most important controls on soil respiration (Conant, Dalla-Betta, Klopatek, & Klopatek, 2004), warming-induced changes in microclimatic variables may alter soil microbial activity, leading to shifts in soil respiration rates (Luo et al., 2001). The lack of consensus on the relative importance of these mechanisms hinders our ability to model long-term soil respiration responses to warming, which are fundamental to increase confidence in soil C projections in a warmer world (Bradford et al., 2016; Zhou et al., 2012).

Beyond heterotrophic microbial CO_2_ production, soil respiration is also a product of plant roots and other autotrophic organisms inhabiting soil surfaces such as biocrusts (topsoil communities formed by cyanobacteria, algae, mosses, liverworts, fungi, bacteria and lichens, Weber, Büdel, & Belnap, 2016). In drylands, which cover 41% of the total land surface (Cherlet et al., 2018) and store 32% of the Earth’s soil organic C (SOC) pool (Plaza et al., 2018), up to 42% of the overall soil respiration comes from biocrust-dominated microsites (Castillo-Monroy, Maestre, Rey, Soliveres, & García-Palacios, 2011; Feng et al., 2014, 2013). Biocrusts are particularly relevant for the global C cycle, as it has been estimated that they cover ca. 12% of the Earth’s terrestrial surface (Rodriguez-Caballero et al., 2018) and fix over 2.6 Pg•yr^−1^ of atmospheric C globally (Elbert et al., 2012). Given their extent and importance for the C cycle, biocrusts are a major ecosystem component when evaluating warming effects on soil respiration in drylands.

Biocrust constituents are severely affected by warming; the physiological performance of soil lichens and mosses have been found to decrease with warming in experiments conducted in Spain, USA, China and South Africa (Grote, Belnap, Housman, & Sparks, 2010; Maestre, Delgado-Baquerizo, Jeffries, Eldridge, & Ochoa, 2015; Maestre et al., 2013; Maphangwa, Musil, Raitt, & Zedda, 2012; Ouyang & Hu, 2017). These responses have been linked to warming-induced reductions in soil moisture and dew events (Ladrón de Guevara et al., 2014; Ouyang & Hu, 2017). Such physiological responses are critical to maintain the photosynthetic activity of biocrust communities (del Prado & Sancho, 2007; Veste, Littmann, Friedrich, & Breckle, 2001), and may lead to dramatic losses in the cover of biocrust-forming lichens and mosses (up to 40%) after a few years of temperature manipulation (Darrouzet-Nardi, Reed, Grote, & Belnap, 2018; Escolar, Martínez, Bowker, & Maestre, 2012; Ferrenberg, Reed, Belnap, & Schlesinger, 2015; Maestre et al., 2013). To evaluate the overall warming effects on soil respiration in drylands, the heterotrophic mechanisms (i.e. substrate depletion, microbial acclimation and changes in microclimatic variables) driving soil respiration responses to warming should be assessed jointly with the shifts in biocrust cover promoted by elevated temperatures (García-Palacios et al., 2018; Maestre et al., 2013).

In drylands, most studies evaluating soil respiration responses to experimental warming have been conducted over short-term periods (Darrouzet-Nardi, Reed, Grote, & Belnap, 2015; Escolar, Maestre, & Rey, 2015; Guan, Li, Zhang, & Li, 2019; Maestre et al., 2013), and consequently long-term warming effects are virtually unknown. To our knowledge, only Darrouzet-Nardi et al., (2018) have explicitly confronted short- vs. long-term soil respiration responses to warming in biocrust-dominated drylands, but no study so far has addressed the heterotrophic and autotrophic mechanisms underlying the transient soil respiration responses to warming. Given the importance of soil carbon-climate feedbacks to forecast greenhouse gas emissions globally (Carey et al., 2016; Crowther et al., 2016), and the extent of drylands worldwide, it is critical to evaluate both short and long-term soil respiration responses to warming in these environments and how these are modulated by biocrusts and soil microbial communities.

Here, we confronted short-term (0-2 years) vs. long-term (8-10 years) soil respiration responses to experimental warming in a biocrust-dominated dryland in central Spain. Data from this experiment were coupled to those from laboratory incubations at four assay temperatures (10, 20, 30 and 40ºC), which allowed us to gain mechanistic insights on the importance of the autotrophic and heterotrophic mechanisms as drivers of soil respiration responses to warming over time. Using this combination of approaches, which to the best of our knowledge has not been used before when evaluating soil respiration responses to warming in drylands, we evaluated: i) short- and long-term warming impacts on soil respiration and its temperature sensitivity (objective i), ii) how warming-induced effects on soil temperature and moisture affect soil respiration responses to elevated temperatures (objective ii), iii) the role of biocrusts as modulators of short- and long-term soil respiration responses to warming (objective iii), and iv) the importance of thermal acclimation of soil microbial respiration as a driver of soil respiration responses to long-term warming (objective iv).

## Materials and methods

### Study area

The study was conducted at the Aranjuez Experimental Station, located in central Spain (40°02′N–3°32′W; elevation = 590 m a. s. l.). Its climate is Mediterranean semiarid, with mean annual temperature and precipitation values (2008-2018 period) of 16,5ºC and 336 mm, respectively. Soils are Gypsiric Leptosol (IUSS Working Group WRB, 2006). Perennial plant coverage is < 40%, and biocrust communities dominated by lichens such as *Diploschistes diacapsis, Squamarina lentigera, Fulgensia subbracteata and Buellia zoharyi*. and mosses *Pleurochaete squarrosa* and *Didymodon acutus* cover ~32% of the soil surface (Castillo-Monroy, Maestre, et al., 2011; Maestre et al., 2013). Cyanobacteria from the genera *Microcoleus, Schizothrix, Tolypothrix, Scytonema* and *Nostoc* also form part of biocrusts at this site (Cano-Díaz, Mateo, Muñoz-Martín, & Maestre, 2018).

In July 2008 we established a full factorial experiment with two factors of two levels each: warming treatment (ambient vs. increased temperature) and initial biocrust cover (low: < 20% vs high: >50%). We installed open top chambers (OTCs) to reach a warming scenario similar to the temperature increase of 2–3°C forecasted for 2040– 2070 in this region in atmosphere–ocean general circulation models (De Castro, Martín-Vide, & Alonso, 2005). OTCs present a hexagonal design made of methacrylate sheets (40 × 50 × 32 cm), a material that according to the manufacturer (Decorplax S.L., Humanes, Spain) ensures 92% transmittance in the visible spectrum and a very low emission in the infrared wavelength. To allow air circulation and so to avoid overheating, OTCs are suspended 3–5 cm over the ground by a metal frame. Ten plots (1.25 x 1.25 m) per combination of treatments were randomly distributed and separated at least 1 m to diminish the risk of lack of independence between replicates (n = 40). We inserted a PVC collar (20 cm diameter, 8 cm height) in each plot to monitor soil respiration and biocrust cover over time. See Escolar et al. (2012) and Maestre et al. (2013) for additional details on the experimental design.

### Testing the warming effects on soil microclimatic conditions

We focused on warming effects on soil temperature and soil moisture as the two main drivers controlling soil respiration in drylands (Castillo-Monroy, Maestre, et al., 2011; Conant et al., 2004; Veste et al., 2001). In parallel to soil respiration measurements, we monitored soil temperature at 0-2 cm depth with protected diodes at the beginning of the experiment and since 2012 (i.e. four years after experimental set-up) with a Li-8100 Automated Soil CO_2_ Flux System (Li-COR, Lincoln, NB, USA) because the later measurements were faster and more accurate. Data obtained with the Li-8100 were corrected using a calibration between both methods (r=0.956, Figure S1). Volumetric soil moisture was measured monthly at 0-5 cm depth using time-domain reflectometry (TDR; Topp & Davis, 1985).

### Soil CO_2_ efflux measurements and its temperature sensitivity (Q10)

The soil CO_2_ efflux rate of the whole soil column, including both biocrusts and soil microbial communities, was measured *in situ* once a month in two contrasted periods: 0-2 yr (short-term hereafter) and 8-10 yr (long-term hereafter) after the setup of the experiment. Measurements were conducted with a closed dynamic system (Li-8100). The opaque chamber used for these measurements had a volume of 4843 cm^3^ and covered an area of 317.8 cm^2^. Given the low CO_2_ efflux rates typically observed in semiarid ecosystems (Castillo-Monroy, Maestre, et al., 2011; Maestre et al., 2013), sampling period was set-up to 120 s to ensure reliable measurements. In every survey, half of the replicates were measured in one day (between 10:00 am and 13:00 pm), and the other half were measured over the next day in the same period. Annual plants were removed from the PVC collars at least 48 hours before soil respiration measurements.

Soil respiration missing data due to technical issues was imputed using the R package missForest (Stekhoven & Bühlmann, 2012) as it was done in similar studies (Darrouzet-Nardi et al., 2015, 2018). The missForest is an iterative imputation algorithm based on random forest models which are considered ensemble-learning methods (Breiman, 2001). The algorithm starts filling the missing data with the variable with the fewest gaps and then iteratively re-fits new imputation models until a stopping criterion is reached. We fit one separated missForest model for each combination of treatments (i.e. four models in total) including soil respiration, temperature and moisture, biocrust cover and sampling date.

We evaluated the temperature sensitivity of soil respiration using Q_10_, defined as the increment in soil respiration when temperature increases by 10ºC and calculated at each plot using the following equations (Luo & Zhou, 2006):

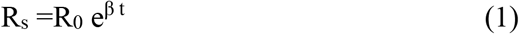

Where R_S_ is soil respiration (μmol m^−2^ s^−1^), R_0_ is the basal soil respiration rate (μmol m^−2^ s^−1^) or intercept of soil respiration at 0ºC, and t is soil temperature (in ºC) measured at the same time as R_S_. β was used to calculate the Q_10_ (increment in R_S_ when t increases by 10 ºC) as follows:

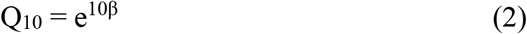

### Monitoring changes in biocrust cover with warming

The total cover of the two major visible components of the biocrust community (lichens and bryophytes) was estimated in each PVC collar at the beginning of the experiment and on a yearly basis hereafter (except during the second year of the experiment, when these measurements were not taken). We used high-resolution photographs to assess the proportion of each collar covered by these biocrust components using the software GIMP (http://www.gimp.org/, to map biocrust-covered areas) and ImageJ (http://rsb.info.nih.gov/ij/, to calculate the size of biocrust-covered areas). Cover estimates obtained with this method are highly related (r^2^=0.84) to those observed in the field with the point sampling method (Ladrón de Guevara et al., 2018). For this study we only considered biocrust surveys included within our sampling periods. Therefore, we used the surveys conducted 1 yr and 9-10 yr (average of both surveys) after the setup of the experiment for the short- and long-term periods, respectively.

### Addressing thermal acclimation of soil microbial respiration using laboratory incubations

We sampled soils (0-5 cm depth) in five field replicates per combination of treatments in 2017, nine years after the setup of the experiment. Biocrust visible components were removed from the samples, which were sieved at 2 mm mesh and stored at 4 ºC for a couple of days until laboratory incubation. We conducted short-term (10 h) laboratory soil incubations at four assay temperatures (10, 20, 30 and 40ºC) as performed in similar mechanistic tests of thermal soil microbial acclimation (Atkin & Tjoelker, 2003; Bradford, Watts, & Davies, 2010; Hochachka & Somero, 2002; Tucker et al., 2013). Soil incubations were performed at 60% of water holding capacity, dark conditions and 100% air humidity.

For each soil sample, we measured soil respiration rates after the addition of two different substrates: sterile deionized water and glucose (at a dose of 10 mg C g^−1^ dry soil) using the MicroResp^TM^ technique (Campbell, Chapman, Cameron, Davidson, & Potts, 2003). The former substrate was used to determine soil basal respiration, whereas the latter was used to account for the effect of substrate limitation on soil respiration rates (Bradford et al., 2010). The glucose dose used in this study is considered to exceed microbial demand (Davidson, Janssens, & Luo, 2006). The MicroResp^TM^ technique (Campbell et al., 2003) is a high-throughput colorimetric method measuring soil respiration rates. We used a CO_2_ detection solution containing cresol red indicator dye that experiences a colour change because of the variation in pH occurring when respired CO_2_ reacts with the bicarbonate of the detection solution. Each microplate well was filled with 150 µl aliquots of the detection solution and was attached to the deep-well microplates containing the soil samples (0.5 g fresh soil/well). Both plates were incubated together at the assay temperature (10, 20, 30 or 40ºC) during the last five hours of the incubation period. The detection plate colour development was read immediately before and after the last five hours of the incubation at 595nm. The colour change in the detection solution was calibrated with an alkali trapping method (*r*^*2*^ =0.86, P <0.001, Lundegardh, 1927).

It is necessary to control for microbial biomass to test for thermal acclimation (Bradford et al., 2010). All available methods to estimate soil microbial biomass have drawbacks (Bradford et al., 2008, 2009; Hartley, Hopkins, Garnett, Sommerkorn, & Wookey, 2009), and hence we measured soil microbial biomass using three different methods to increase the robustness of our results. First, we measured soil induced respiration (µg CO2-C g soil-1 h-1) using autolyzed yeast (Yeast-SIR) as a substrate at 20ºC (Fierer, Schimel, & Holden, 2003). Yeast was added at a dose of 1 mL g soil^−1^ (dry weight equivalent) from a solution containing 12 g of yeast L^−1^ of water. Second, we used a chloroform-fumigation extraction (CFE) (Vance, Brookes, & Jenkinson, 1987). Specifically, we prepared two replicates per sample with almost the same amount of soil: one of the replicates was fumigated with chloroform and the other one remained as a control. Then, we measured total organic carbon (TOC) with an automated TOC analyser in K_2_SO_4_-diluted soil samples. The microbial biomass estimation derived from this technique (mg C kg soil^−1^) was calculated by the difference between fumigated and unfumigated samples. Finally, we measured the relative abundance of soil bacteria (number of DNA copies g-1 soil) using qPCR. The bacterial 16S-rRNA genes were amplified with the Eub 338-Eub 518 primer sets as described in Maestre et al. (2015).

### Statistical analyses

We conducted a series of statistical analyses to achieve the different objectives of the study. To achieve objective i (i.e. how short- and long-term warming affect soil respiration and its temperature sensitivity), we built linear mixed-effect regression models (LMMs) that included warming, initial biocrust cover and their interaction as fixed factors. The temporal dependence of soil respiration measurements across replicates over time (i.e. repeated measures) was tested by including replicate identity and sampling date in the model as random factors. To test the effect of warming on Q_10_, we built linear regression models (LMs) including warming, initial biocrust cover and their interaction as fixed factors.

To achieve objective ii (how warming-induced impacts on soil temperature and moisture affect soil respiration responses to this climate change driver) we first evaluated the effects of warming on soil temperature and moisture using LMMs that included warming, initial biocrust cover and their interaction as fixed factors. Sampling date was included in the model as a random factor. We then calculated the effect size of warming on soil respiration, temperature and moisture for each plot at each period (i.e. short-term and long-term) using the response ratio (RR, Hedges, Gurevitch, & Curtis, 1999):

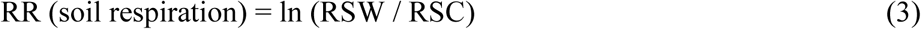

where RSW is the soil respiration in each warmed plot and RSC is the mean soil respiration in the control plots. The RRs were estimated separately for each initial biocrust cover level. To calculate the RRs, we first computed the average across the sampling dates per period for each plot. Then, to test the relationship test the effect of warming-induced changes in soil temperature on the warming effect on soil respiration, we evaluated the relationship between the RR of soil respiration and that of soil temperature using LMs. Similarly, to address the relationship between warming-induces change in soil moisture and soil respiration, we evaluated the relationship between the RR of soil respiration and that of soil moisture using LMs.

To achieve objective iii (the role of biocrusts as modulators of short- and long-term soil respiration responses to warming), we first evaluated the effects of warming on biocrust cover using LMs with warming, initial biocrust cover and their interaction as fixed factors. Then, we evaluated whether warming-induced changes in biocrust cover control soil respiration responses to warming during short- and long-term periods. To do so, we evaluated the relationship between the RR of soil respiration and that of biocrust cover using LMs. The RRs were calculated as described above for soil respiration.

To achieve objective iv (address the importance of thermal acclimation by soil microbial respiration), we tested whether soil heterotrophic microbial respiration acclimates to elevated temperatures after long-term warming. To do so, we statistically controlled for differences in microbial biomass by including it as a covariate in the model (substrate limitation was alleviated in the laboratory incubations using glucose in excess of microbial demand). We used this approach to control for microbial biomass instead of the mass-specific respiration used in previous studies, as ratios are problematic for statistical analyses because they can obscure true relationships among variables (Bradford et al., 2010; Jasienski & Bazzaz, 1999). Specifically, we ran a separate LM for each soil microbial biomass estimation method (i.e. Yeast-SIR, CFE and qPCR). These LMs incorporated warming treatment, initial biocrust cover, the interaction of these two factors, assay temperature and soil microbial biomass as fixed factors, and analysed their effects on potential soil microbial respiration. The interaction between assay temperature and the warming treatment was also tested but removed because it was not significant (p =0.860).

All the statistical analyses were conducted using the R 3.3.2 statistical software (R Core Team, 2015). The LMMs were fit with a Gaussian error distribution using the ‘lmer’ function of the lme4 package (Bates, Mächler, Bolker, & Walker, 2015). All the analyses were performed separately for short-term and long-term sampling periods. Response data were transformed by taking the natural logarithm of each value when needed to meet the assumptions of normality and homogeneity of variance.

## Results

### Short-term and long-term soil respiration and Q_10_ responses to warming

Warming significantly increased soil respiration during the first two years of the experiment in the high biocrust cover plots (Figure 1a, Table S1, p = 0.029). However, these positive effects disappeared in the long-term (i.e. 8 to 10 years after experimental setup; Figure 1b, Table S2, p=0.457). Seasonally, soil respiration rates were consistently greater in autumn and spring, matching major precipitation events over both the short-(Figure 2a) and the long-term (Figure 2b). The Q_10_ was similar in warmed and control plots in the short-term (Figure S2a, p = 0.818), but this variable was a 10% lower (95% CI= 9 to 11%) in warmed than in control plots for both biocrust cover levels in the long-term (Figure S2b, p < 0.001).

**Figure 1.**
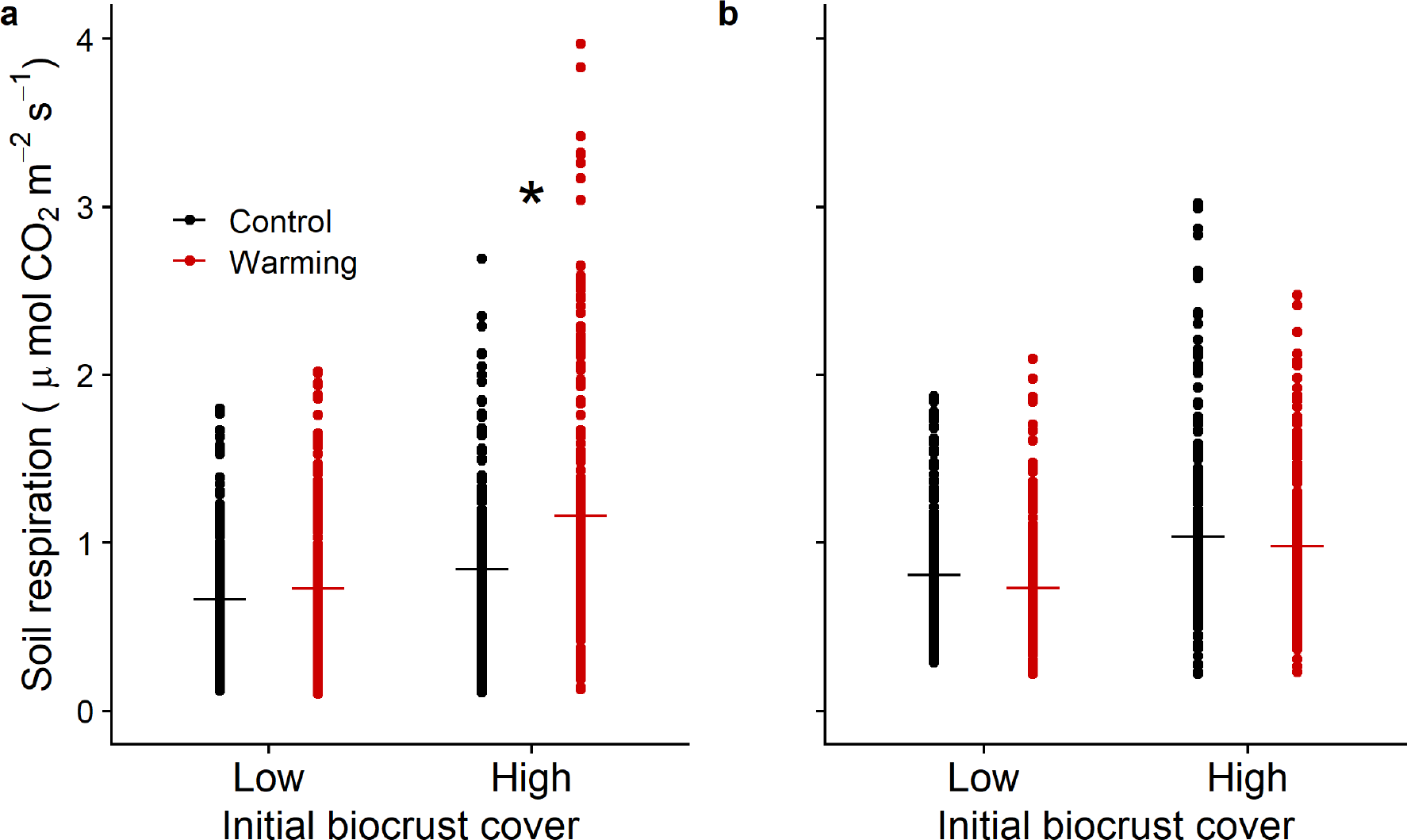
Warming effects on soil respiration rates in the short-term (0 – 2 years after experimental set-up, a) and long-term (8-10 years after experimental set-up, b) at both biocrust cover levels (i.e. low and high). Horizontal lines represent means (n=210 and 240 per combination of treatments, respectively). Asterisks denote significant differences at *p* < 0.05.

**Figure 2.**
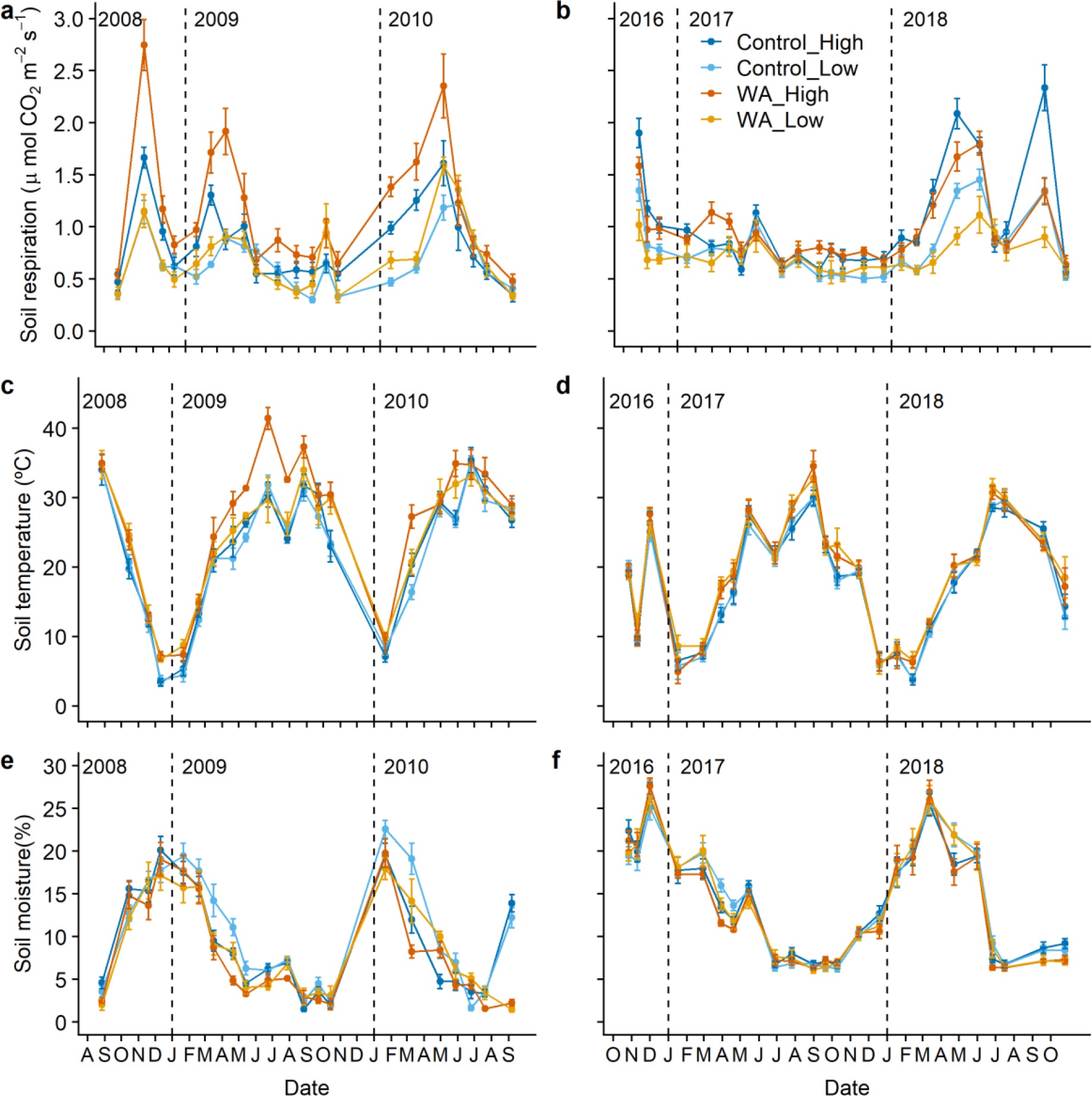
Soil respiration, temperature and moisture measured across short-(after experimental set-up, a, c, e) and long-term (8 – 10 years after experimental set-up, b, d, f) periods. Data are means ± SE (n=10). WA = warming. Low and high refers to initial biocrust cover < 20% and >50%, respectively.

### Changes in soil microclimatic variables as a driver of soil respiration responses to warming

On average, soil temperature was 2.95ºC (95% CI= 2.90 to 2.99 ºC) and 1.43ºC (95% CI= 1.39 to 1.48 ºC) higher in warmed than in control plots at both short and long-term periods, respectively (Figure S3a and b, respectively, p < 0.001 for both periods). In the short-term, mean soil moisture was 1.5% (95% CI= 0.67 to 1.55%) lower in warmed than in control plots (Figure S4a, p< 0.001). However, this effect on soil moisture was not observed under long-term warming (Figure S4b, p=0.227). Seasonally, differences in soil temperature and moisture between control and warmed plots were greater in summer (i.e. from July to September) both in the short-(Figure 2c and e, respectively) and long-term (Figure 2d and f, respectively).

The effect size of warming on soil respiration, as measured with the response ratio, increased when the warming effect on soil temperature was higher under short-term warming (Figure 3a). Contrarily, the effect sizes of warming on soil respiration and soil temperature were not related under long-term warming (Figure 3b). On the other hand, the effect sizes of warming on soil respiration and moisture were not related in the short-(Figure 3c) and long-term (Figure 3d) periods.

**Figure 3.**
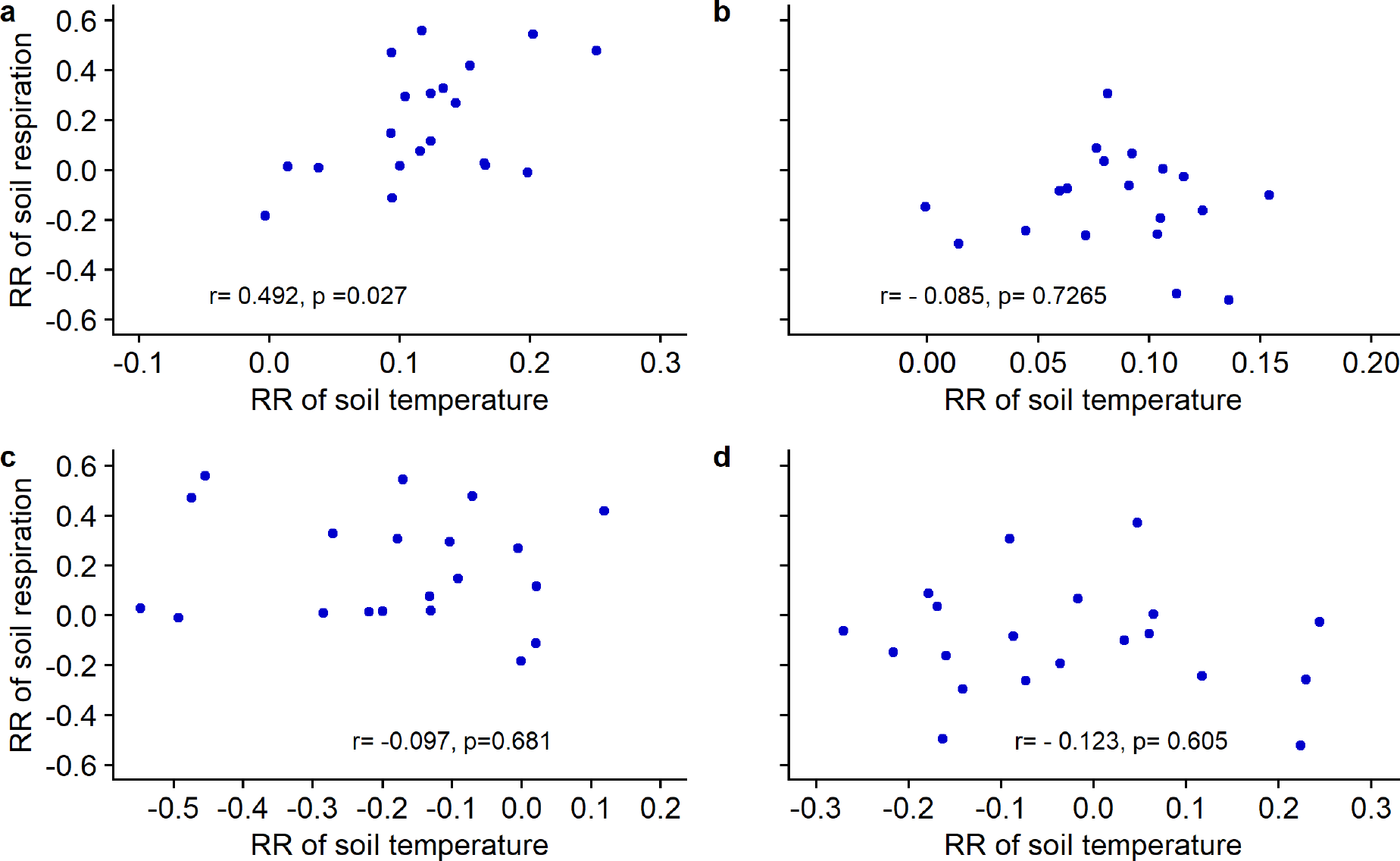
Relationship between the effect size of warming (RR) on soil respiration and on soil temperature in the short-(a) and long-term (b), and between RR of soil respiration and RR of soil moisture in the short-(c) and long-term (d). r refers to the Pearson correlation coefficient and RR are in ln-scale. n = 20 in panels a, c and d and n=19 in panel b.

### Changes in biocrust cover as a driver of soil respiration responses to warming

In the short-term, the total biocrust cover was similar in warmed (9.40%, 95%CI= 8.84 to 9.96% and 66.27%, 95% CI= 63.80 to 68.80%, for low and high initial biocrust cover respectively) and control (7.94%, 95% CI= 7.51 to 8.37% and 64.18%, 95%CI=62.43 to 65.92%, for low and high initial biocrust cover respectively) plots (Figure S5a, p=0.737), when evaluating both initial biocrust cover levels all together. In the long-term, this pattern changed dramatically (Figure S5b), as warming significantly (p< 0.001) decreased total biocrust cover by 26.78% (95% CI= 25.85 to 27.70%) in plots with low initial biocrust cover and by 27.50% (95% CI= 27.17 to 27.83%) in plots with high biocrust cover. The effect size of warming on total biocrust cover and soil respiration were unrelated in the short-term (Figure 4a). However, these effect sizes were significantly and positively related in the longer-term (Figure 4b), indicating that decreases in biocrust cover with warming match with a reduction in soil respiration.

**Figure 4.**
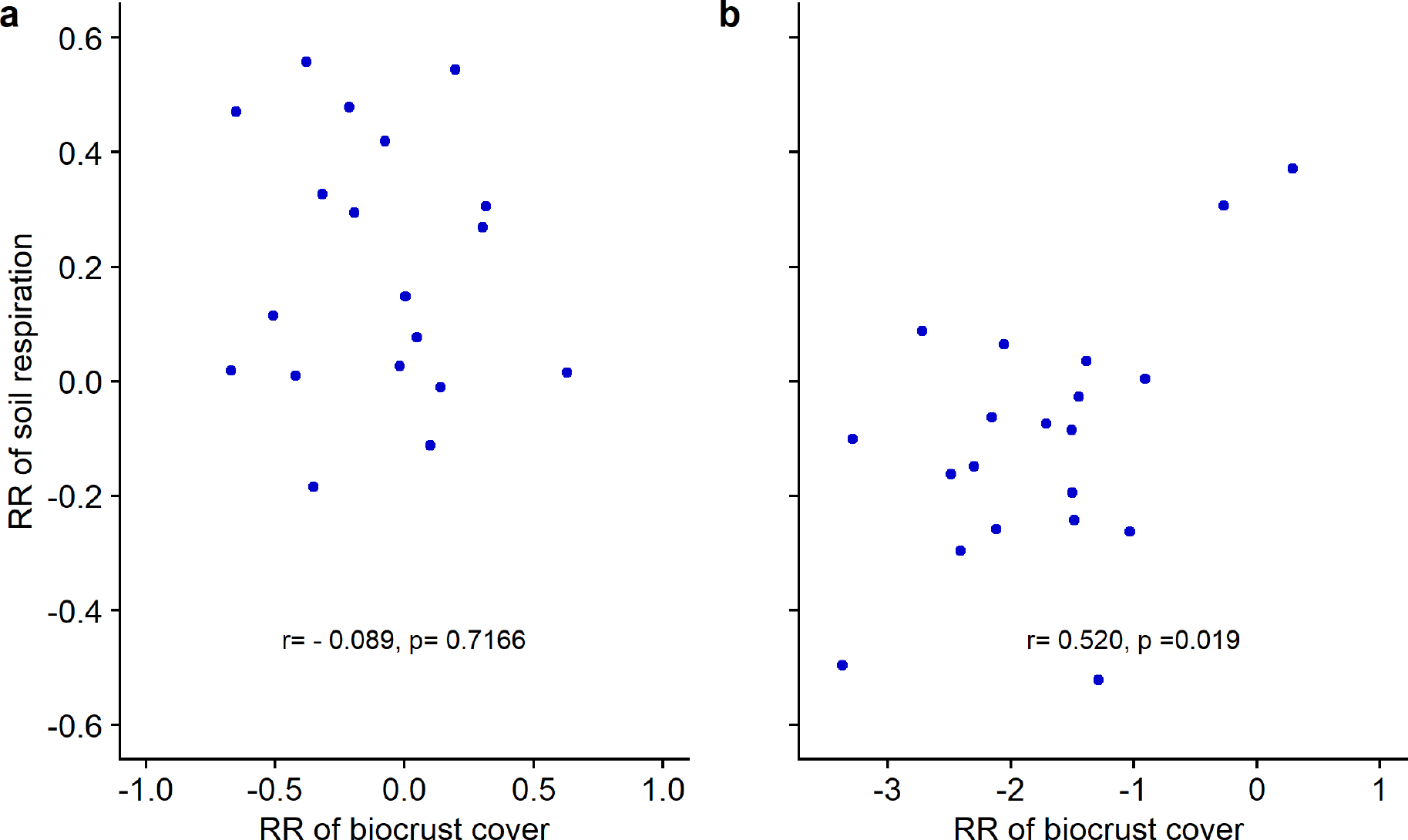
Relationship between the effect size of warming (RR) on soil respiration and on biocrust cover in the short-(a) and long-term (b). r refers to the Pearson correlation coefficient and RR are in ln-scale. n =19 and 20 for panels a and b, respectively.

### Microbial thermal acclimation as a driver of long-term soil respiration responses to warming

Although the positive assay temperature effects on potential soil microbial respiration rates were the largest in magnitude by far (Figure 5, Table S3, p <0.001), we also found a negative effect of warming on soil microbial respiration (Figure 5, Table S3, p = 0.002), which was on average a 30% lower across all assay temperatures. Importantly, this reduction accounted for potential differences in microbial biomass between soil samples (models statistically controlled for differences in microbial biomass), and substrate limitation (incubations were performed with substrate in excess). These results were observed independently of the method used to estimate soil microbial biomass (i.e. Yeast-SIR, CFE or qPCR, Table S3).

**Figure 5.**
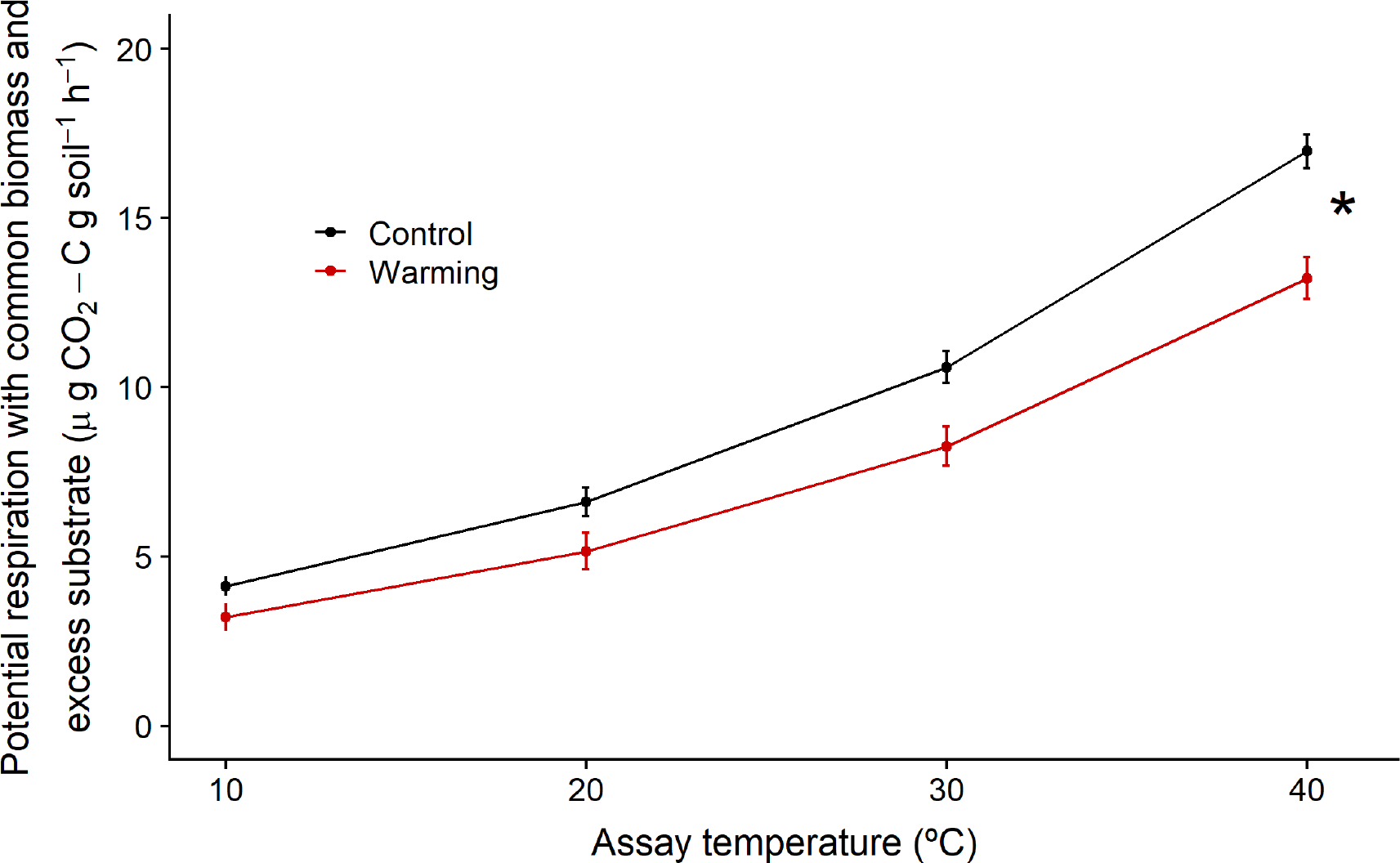
Estimated effects of long-term warming on potential soil respiration rates at a common soil microbial biomass value and with substrate (glucose) in excess of microbial demand. Effect sizes were estimated using coefficients from the ‘Yeast-SIR’ model (Table S3). Specifically, the unstandardized coefficients were used in a regression equation, along with the mean value across all plots for the microbial biomass, one of treatments (i.e. control or warming) and then for assay temperature by systematically increasing the control from the lowest to highest temperatures used in the laboratory incubations. Given that there are not differences between both initial biocrust cover levels, we only represent the data at the low initial biocrust cover. Error bars show the standard deviation. Asterisks denote significant differences at *p* < 0.05.

## Discussion

The positive effect of warming on soil respiration observed in the short-term in plots with high initial biocrust cover disappeared after ten years of warming. This long-term response to warming was linked to a decrease in Q_10_ in the warmed compared to the control plots. Additionally, we found support for several mechanisms driving short and long-term soil respiration responses to warming such as a warming-induced increases in soil temperature, microbial thermal acclimation and changes in total biocrust cover.

Short-term studies have found contrasting soil respiration responses to warming in drylands, ranging from positive (Darrouzet-Nardi et al., 2015; Shen, Reynolds, & Hui, 2009) to negative (García-Palacios et al., 2018; Xu, Hou, Zhang, Liu, & Zhou, 2016). The rare dryland studies that have evaluated warming effects for more than five years have found that soil respiration rates return to control levels after few years of warming (Darrouzet-Nardi et al., 2018; García-Palacios et al., 2018). We compared soil respiration responses to warming in the short vs. long-term and found a positive warming effect in areas with high initial biocrust cover after two years of warming. This positive short-term effect was not sustained through time, and it disappeared after ten years of elevated temperatures. Accordingly, Q_10_ values were significantly lower in the warmed plots compared to the control plots under long-term warming. The mismatch between our short-term results and previous studies also conducted in drylands (García-Palacios et al., 2018; Xu et al., 2016) may be caused by different soil respiration responses to warming due to changes in the mechanisms driving such responses. Such changes in the mechanisms underlying soil respiration responses to warming may also explain the differences between the warming effects on soil respiration observed short-and long-term in our study. Therefore, to better understand soil respiration responses to warming both in the short and long-term, the different drivers that could regulate such responses should be investigated.

Warming-induced increases in soil temperature led to an increase on soil respiration in the short-term, especially in areas with high biocrust cover. Such elevated temperatures effects on soil respiration may be influenced by significant peaks of soil respiration after small rainfall or dew events in the study area (Cable & Huxman, 2004; Ladrón de Guevara et al., 2014). For instance, peaks in soil respiration have been observed in a biocrusted site in the Kalahari Sands (Botswana) after rainfall events of just 1.6 mm (Thomas, Hoon, & Dougill, 2011). Therefore, increases in soil temperature were the main driver underlying the short-term soil respiration responses to warming, given that the mean soil moisture observed (8.5% in the short-term) may be enough to support microbial activity. However, we did not observe this direct effect of warming-induced elevated temperature on soil respiration in the long-term. The disagreement between this result and the expectation that soil respiration should increase with warming (Kirschbaum, 2006) may be a consequence of a long-term effect of the warming treatment on the biocrust and soil microbial communities, compensating the direct effect of increased temperatures. On the hand, experimental warming reduced soil moisture by 1.5% in the short-term, whereas it did not have any effect in the long-term (0.2% reduction). However, the soil respiration responses to warming were independent of changes in soil moisture over both periods. Our results indicate that soil respiration responses to warming are not a product of a reduction in soil microbial activity with warming-induced soil drying, which disagrees with the results found in previous dryland studies (Pendall et al., 2013; Rey et al., 2011). This mismatch between our results and previous findings may be a consequence of the magnitude of the soil moisture change induced by warming. For instance, in Pendall et al., (2013) soil respiration responses to warming were mediated by a 15% decrease in soil moisture.

Therefore, the warming-induced reduction of soil moisture observed in our study may not be large enough to drive soil respiration responses to warming. Additionally, soil respiration in drylands is not only controlled by rainfall events but also by dew generated in the early morning (Rey et al., 2011), as dew-like water inputs were enough to stimulate the respiration of biocrust-forming lichens and the soil microorganisms associated to them (Delgado-Baquerizo, Maestre, Rodríguez, & Gallardo, 2013; Ladrón de Guevara et al., 2014). Therefore, the increased activity of biocrusts, which are a major contributor to soil respiration in our study area, due to water inputs derived from dew events may explain why soil moisture was not driving soil respiration responses to warming neither in the short-nor in the long-term. According to these results, warming-induced changes in soil microclimatic variables do not seem to be the main mechanism underlying soil respiration responses to elevated temperatures in the long-term. Therefore, we tested whether changes in biocrust cover and thermal acclimation of soil microbiota could be the drivers of the soil respiration responses to warming observed.

We observed that soil respiration was larger in the plots with high compared with low initial biocrust cover during both warming periods, albeit temporal trends of soil respiration were similar at both biocrust levels. These results agree with those observed in previous studies showing greater respiration rates in areas with visible and well-developed biocrusts (Castillo-Monroy, Maestre, et al., 2011; Feng et al., 2013). Accordingly, higher soil respiration rates have also found in areas with lichen-dominated biocrusts than in those dominated by mosses or algae (Feng et al., 2014). Therefore, the differences in soil respiration between low and high biocrust cover plots observed in our study may be a result a result of the biological activity of the mosses and lichens directly or through their effect on soil microbial communities (Castillo-Monroy, Bowker, et al., 2011). On the other hand, our results showed that soil respiration responses to short-term warming were independent of changes in biocrust cover, as biocrusts were not affected by the warming treatment in the short-term. Contrary, we observed that soil respiration responses to warming were mediated by warming-induced reduction in biocrust cover in the long-term. The observed decrease in biocrust cover with warming may not seem consistent with previous findings showing that lichens are well adapted to elevated temperatures and are resistant to desiccation (Green, Sancho, & Pintado, 2011). However, it agrees with other studies conducted in drylands which found an important reduction in biocrust cover after some years of warming (Ferrenberg et al., 2015; Maestre, Escolar, et al., 2015). Although clarifying the physiological mechanisms underlying this dramatic decrease in biocrust cover under long-term warming is not a goal of this study, we speculate that it may be due to lichen mortality as a consequence of a reduction of C fixation due to direct and indirect effects of the warming treatment (Ladrón de Guevara et al., 2014). Such reduction, and therefore in the autotrophic soil respiration coming from biocrusts, may explain the decreased soil respiration rates observed in the warmed plots in the long-term. To sum up, our results suggest that biocrusts modulate soil respiration rates and that warming-induced changes in their cover are one of the main drivers governing observing soil respiration responses to long-term warming.

Finally, we found a negative effect of field warming on soil microbial respiration at a common biomass and excess substrate in the laboratory incubations. This result highlights that soil microbial respiration acclimated to warming conditions in this dryland ecosystem, and suggests that thermal acclimation may drive the lack of warming effects on soil respiration over the long-term. Importantly, this result was observed regardless of the method used to measure microbial biomass, suggesting that substrate-induced respiration is an appropriate estimate of microbial biomass to test for thermal acclimation of soil respiration (Bradford et al., 2008, 2009). The negative field warming effect on soil microbial respiration observed is consisted with biochemical acclimation to different thermal regimes reached through evolutionary trade-offs (Hochachka & Somero, 2002). However, we cannot state that biochemical acclimation is the only mechanism operating to explain our results as the ‘aggregate’ soil respiratory activity may be modified by shifts at the individual, population and community levels. Although the observed negative effect of warming on potential soil microbial respiration rates may seem incompatible with the expected positive link between temperature and soil microbial respiration rates (Davidson & Janssens, 2006; Kirschbaum, 2006; Lloyd & Taylor, 1994; Tucker et al., 2013), such positive relationship was supported by the conspicuous positive effect of assay temperature on respiration rates observed in our study. The thermal acclimation of soil respiration observed in this study provides empirical support to previous global extrapolations showing that soil C losses to the atmosphere via soil respiration with elevated temperature may be lower in drylands than in other biomes (Crowther et al., 2016). Indeed, in a global study analysing data from 27 different temperature manipulation experiments, spanning nine biomes, drylands and boreal are the only ecosystems where differences in temperature sensitivity between warmed and control plots have been found (Carey et al., 2016). Therefore, they only found evidence for thermal acclimation of soil respiration in drylands and boreal forests, agreeing with our results.

In conclusion, we found that short-term increases on soil respiration with warming disappeared after ten years of continuous warming in a biocrust-dominated dryland. This pattern was associated with a long-term decreased in temperature sensitivity of soil respiration (Q_10_). Our results suggest that the main driver regulating short-term soil respiration responses to warming was the increase in soil temperature, whereas both thermal acclimation and a dramatic loss of biocrust cover drove soil respiration responses to warming in the long-term. Our results highlight the need to evaluate the effects of warming at both the short- and long-term to better understand soil respiration responses to this climate change driver, and the important role that long-term experiments play for doing so. They also emphasize the need to include both thermal acclimation and biocrust communities in models aiming to forecast soil greenhouse gas emission predictions in drylands, as this would improve our capacity to forecast future temperatures and expand our understanding of C - climate feedbacks.

## Acknowledgements

This research was funded by the European Research Council (ERC Grant agreements 242658 [BIOCOM] and 647038 [BIODESERT]). M.D. is supported by a FPU fellowship from the Spanish Ministry of Education, Culture and Sports (FPU-15/00392). P.G-P. and S.A. acknowledge the Spanish MINECO for financial support via the DIGGING_DEEPER project through the 2015-2016 BiodivERsA3/FACCE-JPI joint call for research proposals. F.T.M. and S.A. acknowledge support from the Generalitat Valenciana (CIDEGENT/2018/041).

## Authorship

F.T.M. designed the field study and wrote the grant that funded the work. F.T.M, P.G.P and M.D. developed the original idea of the analyses presented in the manuscript. M.D. performed the statistical analyses, with inputs from F.T.M and P.G.P. M. D., S.A., B.G. and V. O. conducted the field and laboratory work. All authors contributed to data interpretation. M.D. wrote the first version of the manuscript, which was revised by all co-authors.

