## Supplementary Figure for "Abiotic and biotic drivers underly short- and long-term soil respiration responses to experimental warming in a dryland ecosystem"

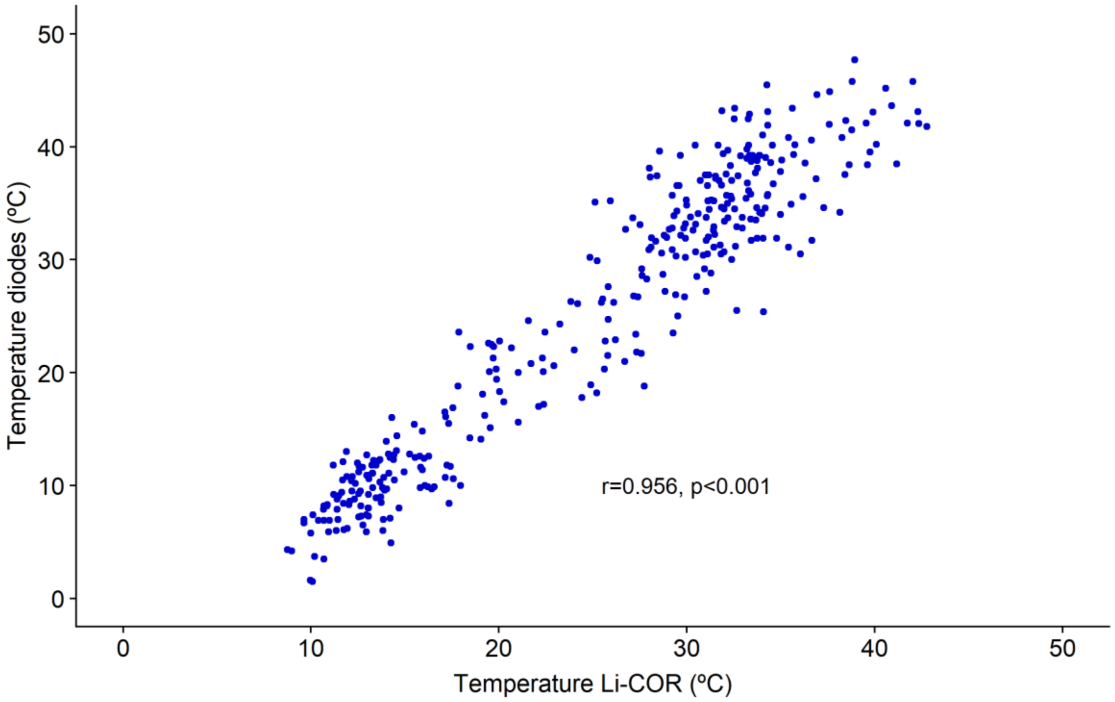


**Figure S1.** Calibration between the two methods used to measure soil temperature throughout the experiment (soil diodes and Li-COR 8100). n=350.

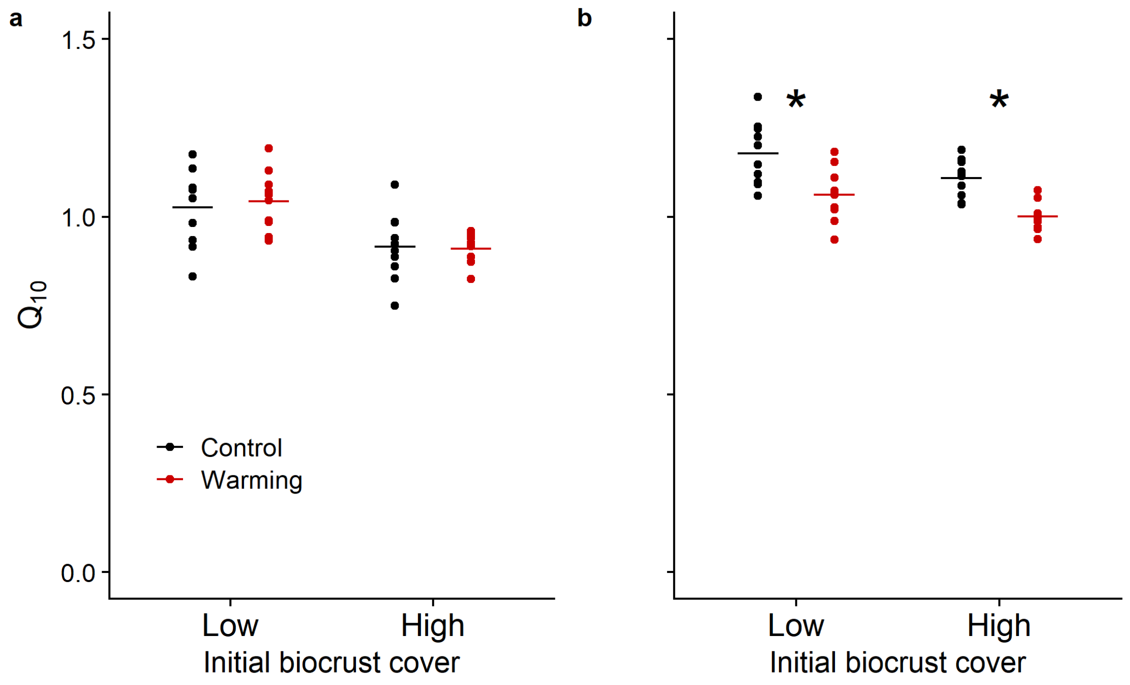


**Figure S2.** Warming effects on the temperature sensitivity of soil respiration ($Q_{10}$) in the short- (0 – 2 years of continuous warming, a) and the long-term (8- 10 years of continuous warming, b) at both initial biocrust cover levels. Horizontal lines represent means (n=10 per combination of treatments). Asterisks denote significant differences at $p < 0.05$.

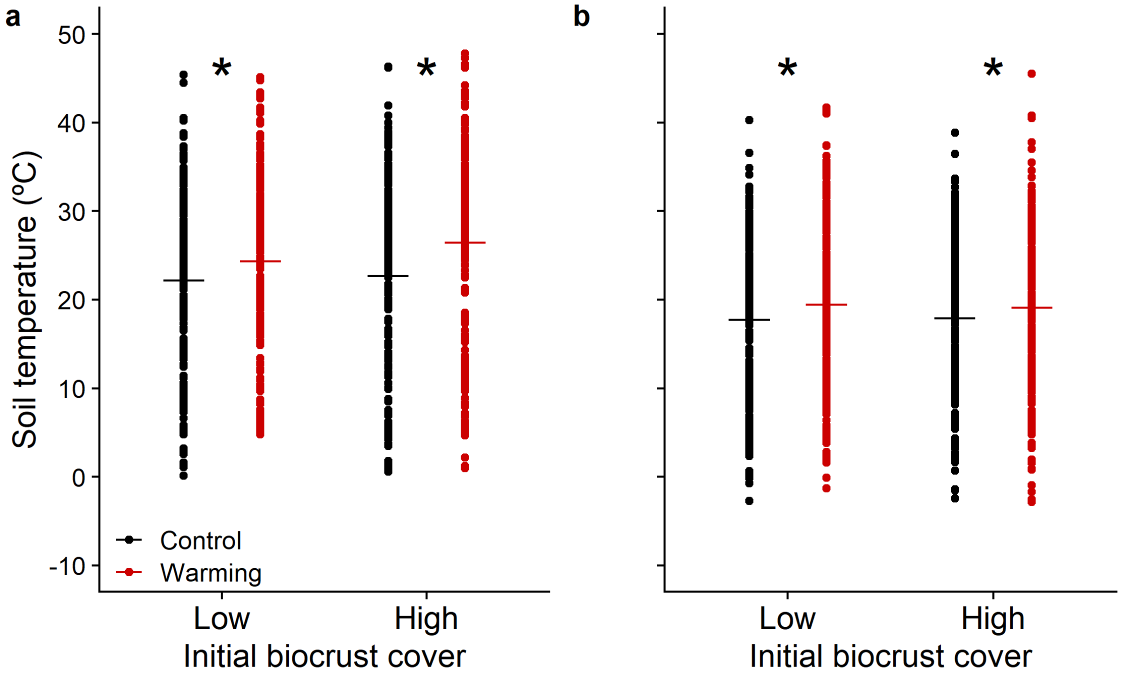


**Figure S3.** Warming effects on soil temperature in the short- (0 – 2 years of continuous warming, a) and the long-term (8- 10 years of continuous warming, b) at both initial biocrust cover levels. Horizontal lines represent means (n=210 and 240 per combination of treatments, respectively). Asterisks denote significant differences at $p < 0.05$.

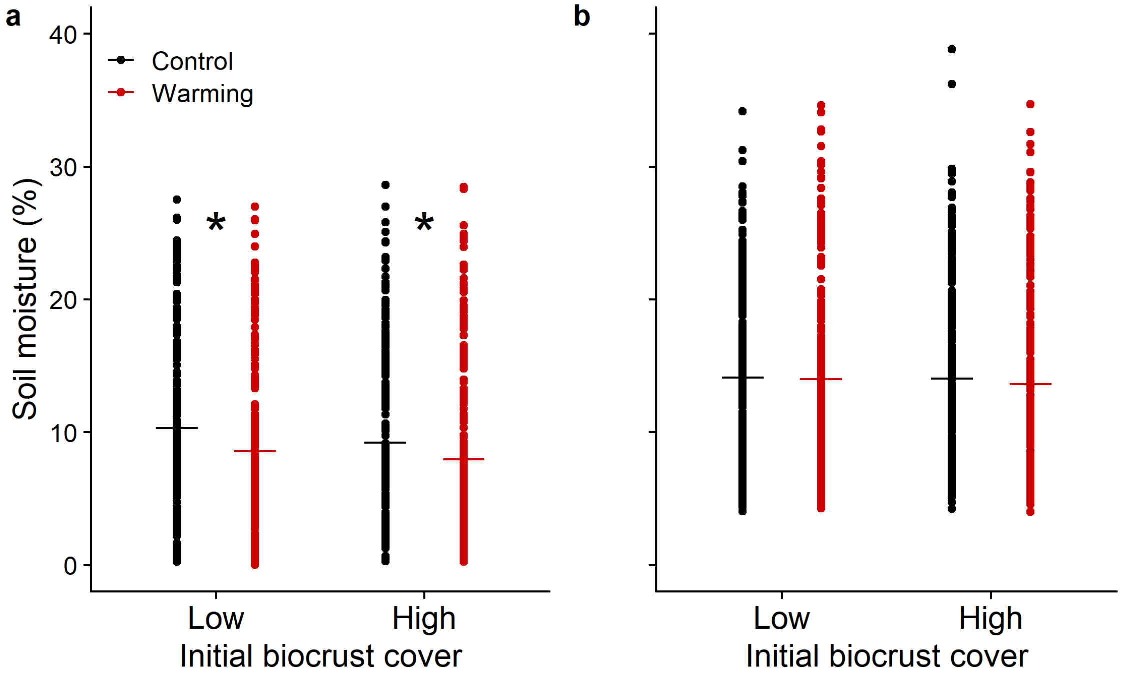


**Figure S4.** Warming effects on soil moisture in the short- (0 – 2 years of continuous warming, a) and the long-term (8- 10 years of continuous warming, b) at both initial biocrust cover levels. Horizontal lines represent means (n=210 and 240 per combination of treatments, respectively). Asterisks denote significant differences at $p < 0.05$.

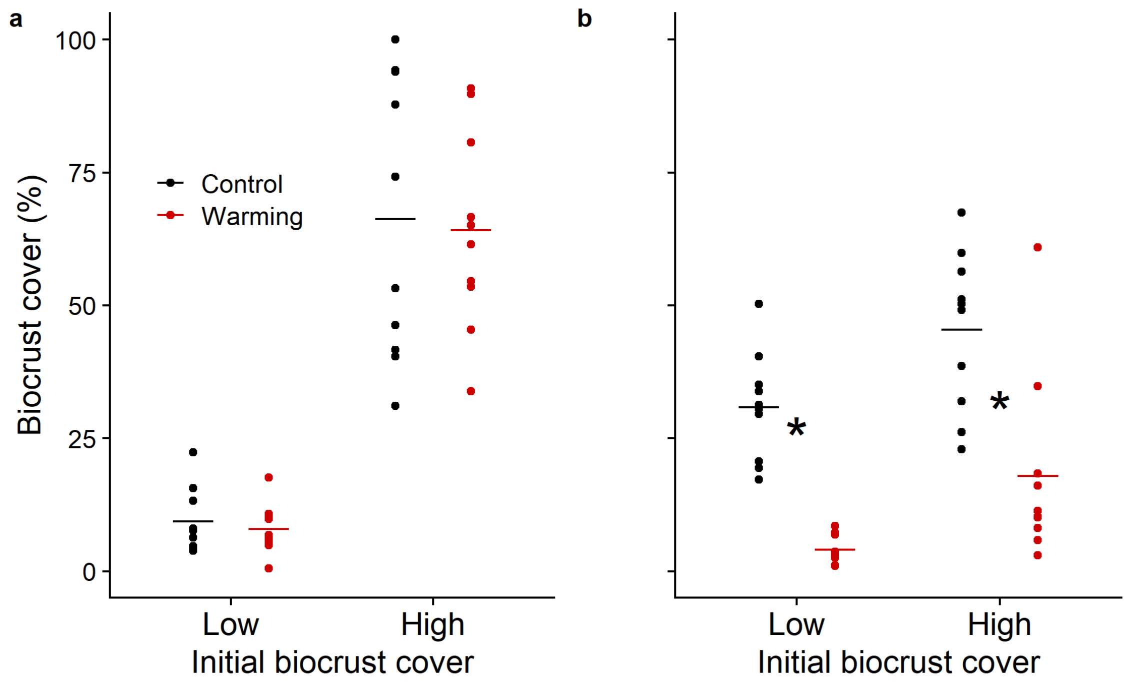


**Figure S5.** Total biocrust cover throughout the duration of the experiment in the short- (0 – 2 years of continuous warming, a) and the long-term (8- 10 years of continuous warming, b) at both initial biocrust cover levels. Horizontal lines represent means (n=10 per combination of treatments). Asterisks denote significant differences at $p < 0.05$.

**Table S1.** Summary results of the linear mixed-effect regression models (LMMs) and ANOVA table used to test short-term soil respiration response to warming. The table shows the unstandardized coefficients (mean ± SD) of the model. Respiration rates were ln-transformed to meet normality assumptions. We used 10 replicates per combination of treatments and 21 sampling dates (total n = 840). P values below 0.05 are shown in bold.

| Variables | Coefficients | df | F | P |
|---|---|---|---|---|
| Intercept | - 0.536 ± 0.107 | 34 | - | **< 0.001** |
| Warming | 0.037 ± 0.081 | 36 | 8.545 | **0.006** |
| Biocrust cover | 0.189 ± 0.081 | 36 | 31.143 | **< 0.001** |
| Warming x Biocrust | 0.260 ± 0.114 | 36 | 5.178 | **0.029** |

**Table S2.** Summary results of the linear mixed-effect regression models (LMMs) and ANOVA table used to test long-term soil respiration response to warming. The table shows the unstandardized coefficients (mean ± SD) of the model. Respiration rates were ln-transformed to meet normality assumptions. We used 10 replicates per combination of treatments and 24 sampling dates (total n = 960). P values below 0.05 are shown in bold.

| Variables | Coefficients | df | F | P |
|---|---|---|---|---|
| Intercept | - 0.298 ± 0.084 | 32 | - | **0.002** |
| Warming | -0.104 ± 0.082 | 36 | 1.062 | 0.310 |
| Biocrust cover | 0.211 ± 0.082 | 36 | 19.199 | **< 0.001** |
| Warming x Biocrust | 0.087 ± 0.116 | 36 | 0.565 | 0.457 |

**Table S3.** Coefficients (mean ± SD) values for the linear mixed-effects models used to assess long-term thermal adaptation of potential soil microbial respiration (measured with substrate in excess). The table shows the unstandardized coefficients of the Yeast-SIR model and the standardized of all the models. Respiration rates were ln-transformed to meet normality assumptions. Unstandardized coefficients of the Yeast-SIR model were used when plotting Figure 5. We used five replicates per combination of treatments (total n = 80). Coefficients with an associated P < 0.05 are shown in bold.

| Model | | | | |
|---|---|---|---|---|
| **Variables** | **Unstandardized coefficients** | **Standardized coefficients** | | |
| | **Yeast-SIR** | **Yeast-SIR** | **CFE** | **qPCR** |
| Intercept | **0.995 ± 0.191** | **1.964 ± 0.042** | **1.964 ± 0.042** | **1.964 ± 0.042** |
| Assay temperature | **0.047 ± 0.004** | **1.060 ± 0.084** | **1.061 ± 0.084** | **1.061 ± 0.084** |
| Warming | **- 0.250 ± 0.119** | **-0.277 ± 0.084** | **-0.275 ± 0.084** | **-0.277 ± 0.084** |
| Biocrust cover | - 0.046 ± 0.120 | -0.072 ± 0.084 | -0.081 ± 0.084 | -0.070 ± 0.084 |
| Warming x Biocrust | -0.013 ± 0.168 | -0.053 ± 0.168 | -0.052 ± 0.167 | -0.048 ± 0.167 |
| Microbial biomass | 0.034 ± 0.042 | -0.026 ± 0.085 | 0.043 ± 0.085 | -0.016 ± 0.091 |
